# The MicroRNA pathway regulates obligatory aestivation in a flea beetle

**DOI:** 10.1101/2025.02.05.631883

**Authors:** Gözde Güney, Kerstin Schmitt, Johan Zicola, Umut Toprak, Michael Rostás, Stefan Scholten, Doga Cedden

**Affiliations:** Agricultural Entomology, Department of Crop Sciences, University of Göttingen, Göttingen, Germany; Institute of Microbiology and Genetics, Department of Molecular Microbiology and Genetics, University of Göttingen, Göttingen, Germany; Göttingen Center for Molecular Biosciences (GZMB), Service Unit LCMS Protein Analytics, University of Göttingen, Göttingen, Germany; Division of Crop Plant Genetics, Department of Crop Sciences, University of Göttingen, Göttingen, German; Ankara University, Faculty of Agriculture, Department of Plant Protection, Ankara, Turkey; Department of Evolutionary Developmental Genetics, University of Göttingen, Johann-Friedrich-Blumenbach Institute, Göttingen Center for Molecular Biosciences, Göttingen, Germany

**Author notes:** These two authors have contributed equally to this work (Gözde Güney, Doga Cedden). Doga Cedden, Stefan Scholten, Michael Rostás **Email:**.

**Keywords:** gene regulation, aestivation, miRNA, diapause, *Psylliodes chrysocephala*

## Abstract

Aestivation is a dormant state that allows animals to withstand hot and dry summer conditions and requires complex gene regulation. Nevertheless, the mechanisms involved in the regulation of genes necessary for aestivation remain unclear. MicroRNA (miRNA) are known to fine-tune gene expression at the post-transcriptional level and are important for various biological processes. In this study, we investigated the role of the miRNA pathway in the regulation of the obligatory aestivation stage in the cabbage stem flea beetle, a major pest of oilseed rape. Small RNA sequencing showed that ∼25% of miRNAs were differentially abundant during aestivation. The inhibition of the miRNA pathway deregulated 116 proteins in aestivation, which were mainly associated with metabolism and catabolism, including peroxisome activity. Most proteins regulated by miRNA exhibited lower transcript levels during aestivation. RNA degradome sequencing confirmed the miRNA-mediated exonucleolytic decay of several transcripts. Furthermore, inhibiting the miRNA pathway resulted in altered body composition, compromised metabolic suppression, and lower resilience to high temperature during aestivation. Also, beetles could not suppress their feeding activity during the transition into aestivation. Our findings highlight the critical role of miRNA in regulating aestivation in the cabbage stem flea beetle, with important implications for climate change.

## Introduction

Most biological phenomena depend on specific gene expression patterns that are regulated by epigenetic mechanisms (1). The microRNA (miRNA) pathway has been recognized as a key post-transcriptional regulator that enables intricate control of gene expression, complementing canonical epigenetic mechanisms such as histone modifications (2, 3). In animals, the miRNA pathway begins with the transcription of the primary (pri)-miRNA, a 5’-capped and 3’ polyadenylated transcript, by RNA polymerase II in the nucleus. Pri-miRNA is processed by the enzyme Drosha into pre-miRNA hairpins (∼100 nt), which are then transported into the cytoplasm via Exportin-5 (4, 5). In the cytoplasm, the enzyme Dicer-1 further processes pre-miRNA, releasing the miRNA/miRNA* duplex (∼22 nucleotides) (6, 7). One of the strands, known as the mature miRNA, is retained by the Argonaute-1 (Ago-1), forming the RNA-induced silencing complex (RISC) (8). miRNA-loaded-RISC mainly interacts with 3’ untranslated regions (UTR) of mRNA that are recognized mainly by the seed sequence (2-8 nucleotides from the 5’ end) of the miRNA, resulting in decay or translational inhibition of target mRNAs in animals (9).

The miRNA pathway has been implicated in the regulation of various biological processes in animals, including development and metabolism (10–13). Previous studies in insects have also suggested that miRNA could regulate diapause, which is a state of dormancy enabling insects to survive harsh seasonal conditions such as cold winter and hot summer (14–16). Diapause in insects can be referred to as hibernation if it coincides with winter and as aestivation if it coincides with summer (17). MiRNA-mediated regulation of diapause is plausible because this dormant state necessitates the downregulation of numerous genes, a process that can be mediated by the expression of specific miRNAs. However, in non-model organisms, molecular methods are mostly limited to studying miRNA that are differentially abundant between diapausing and non-diapausing stages through small RNA sequencing (14–16). Moreover, past work in this area is highly restricted to *in silico* prediction of miRNA-target genes, which is known to be not very accurate, rather than experimental evidence (18, 19). Furthermore, functional validation of miRNA-mediated regulation of the diapause phenotype is currently lacking. Hence, we lack concrete evidence for miRNA-mediated regulation of diapause-associated genes, and it is unclear whether this regulation is significant at the phenotypic level.

The cabbage stem flea beetle (CSFB), *Psylliodes chrysocephala* (Coleoptera: Chrysomelidae), is a key pest of winter oilseed rape (*Brassica napus napus*) in Northern Europe (20, 21). CSFB eggs hatch in late autumn, and larvae feed by tunneling into the petioles and stems of oilseed rape. After overwintering as larvae, they pupate in the soil in late spring and adults emerge in early summer and feed on oilseed rape leaves (22–24). CSFB adults exhibit obligatory aestivation, meaning that regardless of environmental and rearing conditions, newly emerged adults enter a preparatory (pre-aestivation) phase and, within 15 days, enter aestivation, which lasts from mid to late summer in nature (25). Aestivating CSFB has dramatically reduced metabolic rate and feeding activity, and instead, it relies on internal energy reserves (e.g., lipid stores) to maintain its suppressed metabolism. During aestivation, CSFB demonstrates resilience to elevated temperatures and reduced humidity, conditions characteristic of European summers (25). Only post-aestivation CSFB can reproduce, making aestivation a key stage from a pest management perspective.

In a previous study, we identified thousands of differentially abundant transcripts in aestivation compared to non-aestivation stages, which were associated with key aestivation phenotypes, including lower metabolic rate (25). Yet the mechanisms underlying this gene regulation remained uncharacterized. In this study, we hypothesized that a subset of genes that are required to be suppressed during aestivation are regulated via the miRNA pathway, as this pathway is known to regulate many biological processes through gene suppression. To that end, we first sequenced small RNAs isolated from the three adult stages of CSFB and identified differentially abundant miRNA in aestivation. RNA degradome was sequenced to find evidence for transcript decay mediated by differentially abundant miRNA. In parallel, we knocked down the expression of two core miRNA pathway genes via RNA interference to identify deregulated proteins and the changes in the aestivation phenotype, including resilience to summer conditions and metabolic alterations.

## Results

### Changes in the abundance of miRNA in aestivation

CSFB obligatory enters aestivation within 2 weeks following adult eclosion, dividing adulthood into three stages: pre-aestivation (first 2 weeks), aestivation (15- to 45-day-old adults), and post-aestivation (older than 45-day-old adults). We conducted small RNA-seq in these three stages to *de novo* identify miRNA and investigate differentially abundant miRNA. In total, we identified 105 mature miRNA in CSFB adults based on mirdeep2 scores (Dataset S1), which take into account mature and star miRNA counts and randfold value for the miRNA precursor (an example is provided in Fig. 1A), and 69 of these showed homology to miRNA identified in red flour beetle *Tribolium castaneum* (https://www.mirbase.org/). Due to the absence of a well-established miRNA database for beetles, the miRNAs were named based on their genomic loci of origin.

**Figure 1.**
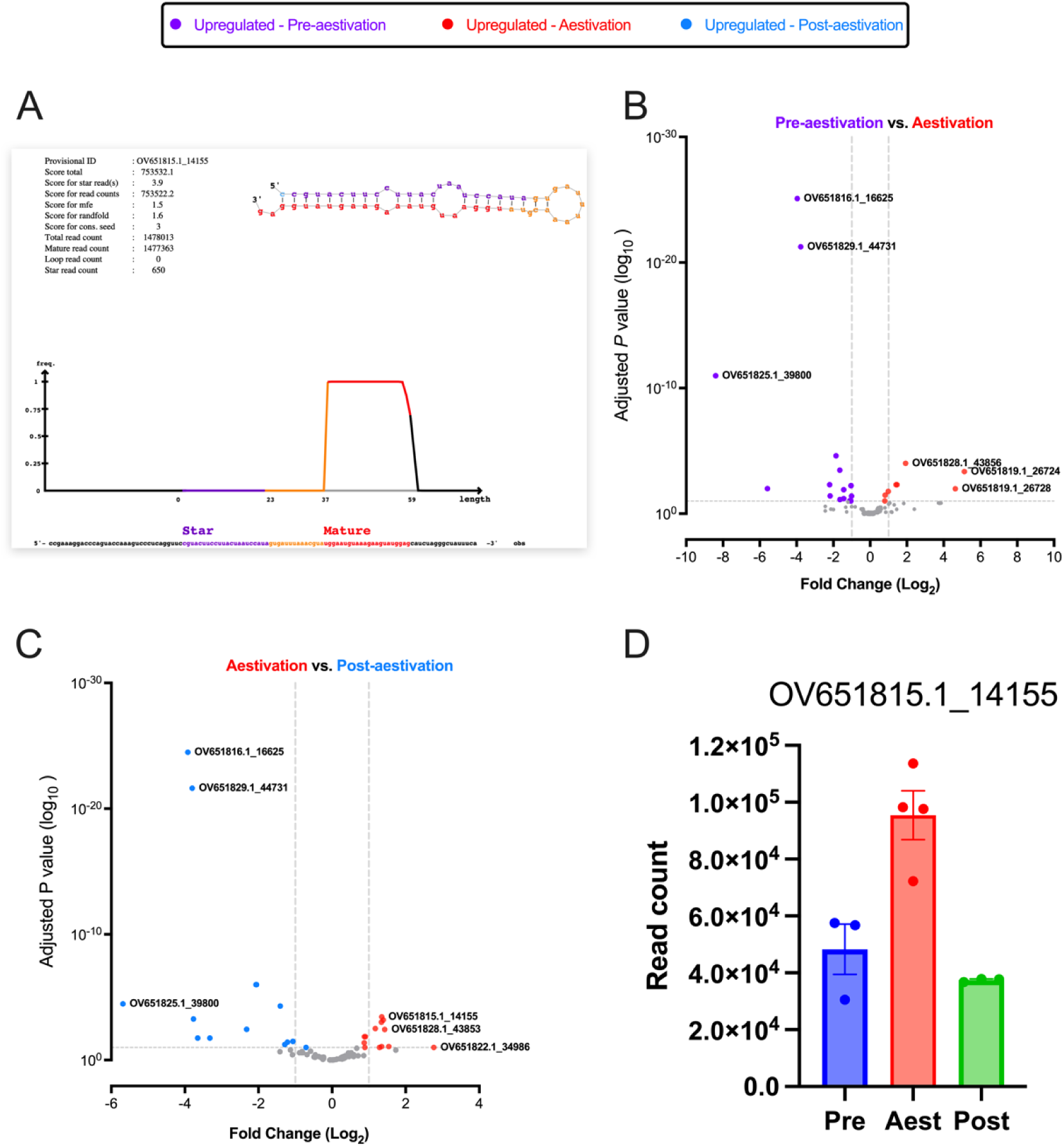
Differentially abundant miRNA at different life stages. A) An example plot produced using miRdeep2 for the identification of a miRNA, which was highly expressed in aestivation compared to other stages. The predicted RNA structure of the pre-miRNA and frequency of read counts corresponding to the mature (red), loop (orange), and passenger (purple) parts of the duplex are shown. DESeq2 analysis was conducted on miRNA identified and quantified using miRdeep2 in pre-aestivation (n = 3), aestivation (n = 4) and post-aestivation (n = 3) cabbage stem flea beetle females. B) Differentially abundant miRNA in the comparison between aestivation and pre-aestivation. C) Differentially abundant miRNA in the comparison between aestivation and post-aestivation. D) Normalized expression count (mean ± SEM) of the example miRNA at three different life stages; pre-aestivation (pre), aestivation (aest), and post-aestivation (post).

In the pre-aestivation vs. aestivation comparison, we identified 8 and 17 miRNAs with significantly higher and lower abundance in aestivation, respectively (adjusted P < 0.05 by DeSeq2, Fig. 1B). The post-aestivation vs. aestivation comparison identified 13 and 15 miRNAs with significantly higher and lower abundance in aestivation, respectively (adjusted P < 0.05 by DeSeq2, Fig. 1C). Four miRNAs had significantly higher abundance in aestivation compared to both pre- and post-aestivation stages (an example is provided in Fig. 1D). Interestingly, we observed that the precursors of two miRNA increasing in abundance in aestivation compared to both other stages mapped 24.7 kb apart on chromosome 16 (Accession: OV651828.1), suggesting that the expression of these two miRNAs might be co-regulated. These two miRNAs were homologous to tca-miR-34-5p and tca-miR-277-3p. Overall, the small RNA sequencing results suggest that a subset of miRNA (∼25% of all miRNA identified in adults) is differentially abundant depending on the life stage, including the aestivation.

### Inhibiting miRNA pathway changes protein levels in aestivation

The differential abundance of miRNA in aestivation suggested that they are important for fine-tuning gene expression to enable aestivation. Hence, we inhibited the miRNA pathway using a chimeric dsRNA that targeted both *dicer-1* and *drosha* (dsDcr1/Dro), which are two key nucleases in miRNA biogenesis. This approach was used to ensure the simultaneous inhibition of both important steps in miRNA biogenesis. We fed CSFB adults with dsDcr1/Dro or dsmGFP (control) upon adult emergence throughout the pre-aestivation period. We confirmed through RT-qPCR that both target genes were significantly knocked down on 15 days post-emergence (i.e., early aestivation) in the dsDcr1/Dro-fed CSFB compared to the control CSFB (*dicer-1:* P < 0.001, t = 7.9 and *drosha*: P < 0.001, t = 6.1 by Šídák’s test, df = 8, Fig. 2A). Furthermore, the proportion of miRNAs within total small RNAs was significantly reduced by 31.9% at 15 days post-emergence in dsDcr-1/Dro-fed CSFB compared to the control CSFB (P = 0.04, t = 2.2, df = 4 by t-test, Fig. 2B).

**Figure 2:**
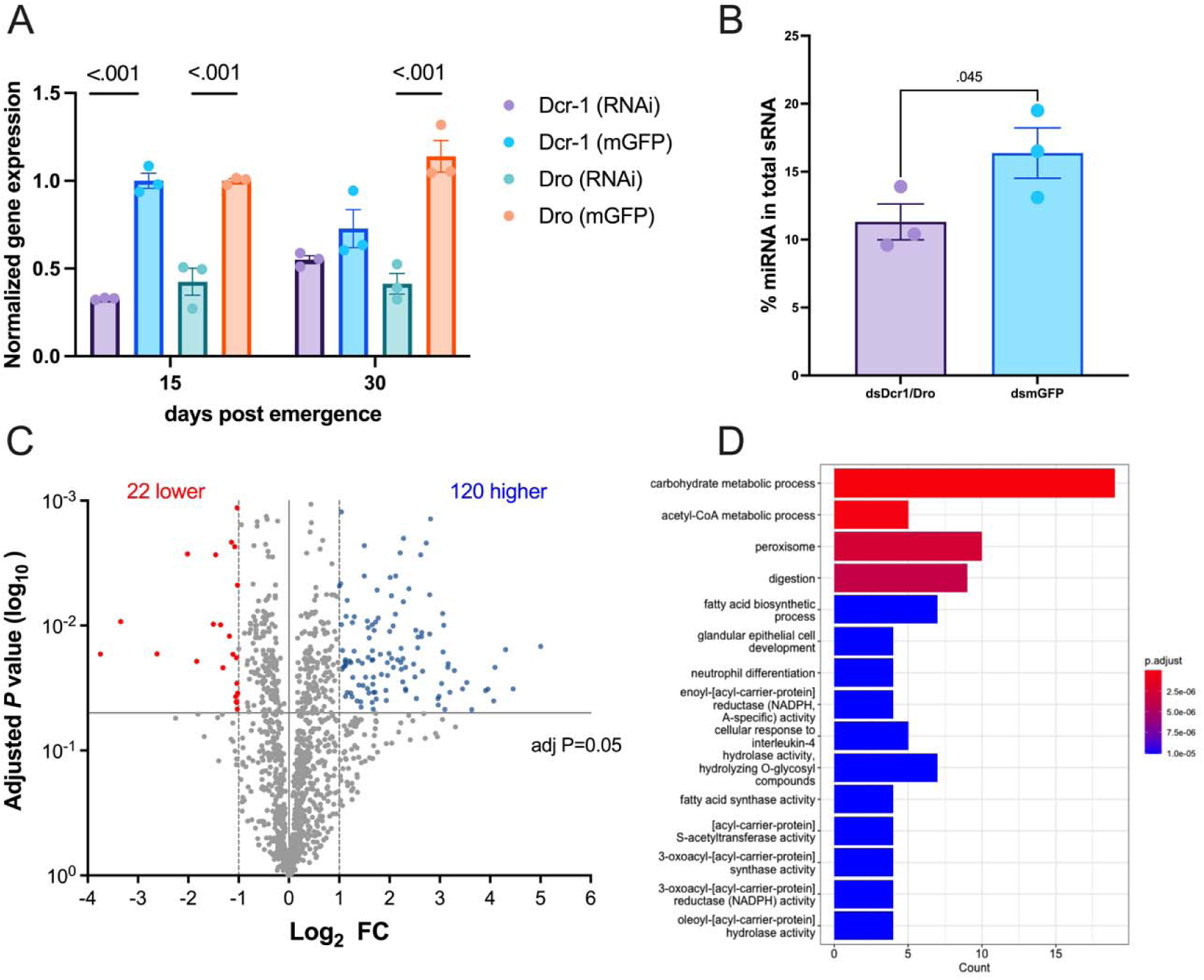
Changes at protein levels following inhibition of miRNA pathway in aestivation. A) Normalized gene expressions of Pc-dicer-1 and Pc-drosha upon the feeding of chimeric dsDcr1/Dro, which simultaneously targeted both genes or control dsRNA mGFP as control (n = 3, mean ± SEM, normalized to the respective gene expression in mGFP group). B) Percent miRNA in total sRNA measured through fragment analysis (n = 3, mean ± SEM). C) Differentially abundant proteins in dsDcr1/Dro fed beetles in comparison to dsmGFP fed beetles (n = 5). Proteins with Log2 FC higher than 1 or lower than −1 in addition to adjusted P below 0.05 were accepted as significantly increased or decreased in abundance. The differential abundance analysis on proteins quantified through LC/MS was conducted using Perseus (v1.4). D) The proteins significantly increased in abundance in dsDcr1/Dro fed beetles were subjected to Gene Ontology (GO) enrichment analysis using clusterProfiler (v4.1). GO terms with P < 0.05 were accepted as significantly enriched.

Next, we assessed the protein levels in aestivating CSFB. In total, our proteomics approach was able to capture 2216 proteins with sufficient abundance for quantification in CSFB adults. We identified 116 proteins with significantly higher abundance in dsDcr1/Dro-fed aestivating CSFB compared to control aestivating CSFB (adjusted P < 0.05 by Welch’s test, Fig. 2C, Dataset S2). We assumed that all 116 proteins were directly regulated by miRNA; however, it is also possible that regulation of a subset of these proteins occurs indirectly, for example, through the regulation of an intermediate transcription factor. In contrast, 22 proteins had lower abundance in dsDcr1/Dro fed aestivating CSFB (adjusted P < 0.05 by Welch’s test). Such a ratio—five times more proteins with higher abundance in dsDcr1/Dro fed aestivating CSFB compared to control aestivating CSFB—was expected because reduction in miRNA-mediated repression should increase levels of proteins normally repressed by miRNA. In contrast, the proteins that decrease in abundance either reflect rare cases where miRNA positively regulates a protein or represent an indirect effect, such as compensatory regulation due to the inhibition of the miRNA pathway. Among the 116 proteins showing higher abundance in dsDcr1/Dro fed CSFB, more than half (73 proteins, 61%) also showed a significant decrease in abundance in aestivation compared to the pre-aestivation in the control CSFB (Fig. S1). This overlap shows that our RNAi approach mostly inhibited the suppression (i.e., increased the abundance) of proteins that are suppressed for aestivation in intact CSFB. In contrast, we observed no overlap between proteins with lower abundance in control aestivating CSFB compared to pre-aestivation control CSFB and those with lower abundance in dsDcr1/Dro-fed aestivating CSFB compared to control aestivating CSFB. In other words, none of the proteins downregulated during aestivation in intact beetles showed reduced levels upon the inhibition of the miRNA pathway. This finding highlights the specificity of RNAi-mediated suppression of the miRNA pathway, combined with proteomics, in identifying miRNA-regulated proteins. Overall, these results suggest that the inhibition of the miRNA pathway mostly deregulated proteins that are suppressed during aestivation, which enabled us to further investigate the role of miRNA in aestivation.

The GO enrichment analysis on proteins with higher abundance in dsDcr1/Dro-fed aestivating CSFB compared to the control aestivating CSFB showed that the deregulated proteins were mainly related to metabolic pathways, including carbohydrate and lipid metabolism and peroxisome activity (Fig. 2D, see Dataset S2 for details). Furthermore, a catabolism related term, namely digestion, was also among the most enriched pathways in the proteins increasing in abundance in dsDcr1/Dro fed aestivating CSFB. These terms suggest that miRNA may have an impact on various aspects of the aestivation phenotype, including energy storage (e.g., lipid reserves), metabolic rate, and feeding.

### Characterizing the mode of regulation by miRNA in aestivation

Following the identification of miRNA-suppressed proteins in aestivation, we sought to characterize the mode of regulation by miRNA. Transcripts encoding 82 out of the 116 proteins showing higher abundance in dsDcr1/Dro fed aestivating CSFB also had lower abundance in aestivation compared to the pre-aestivation (Fig. 3A-B, RNA-seq data from (25)). This suggests that most (71%) of the proteins suppressed through miRNA in aestivation were regulated at the mRNA level. Nonetheless, transcripts encoding for 38 proteins with higher abundance in dsDcr1/Dro fed aestivating CSFB did not change in abundance in aestivation, suggesting these proteins are regulated via translational inhibition, without RNA decay. Alternatively, these 38 proteins might be regulated at the protein level such as decrease in turn-over rate.

**Figure 3:**
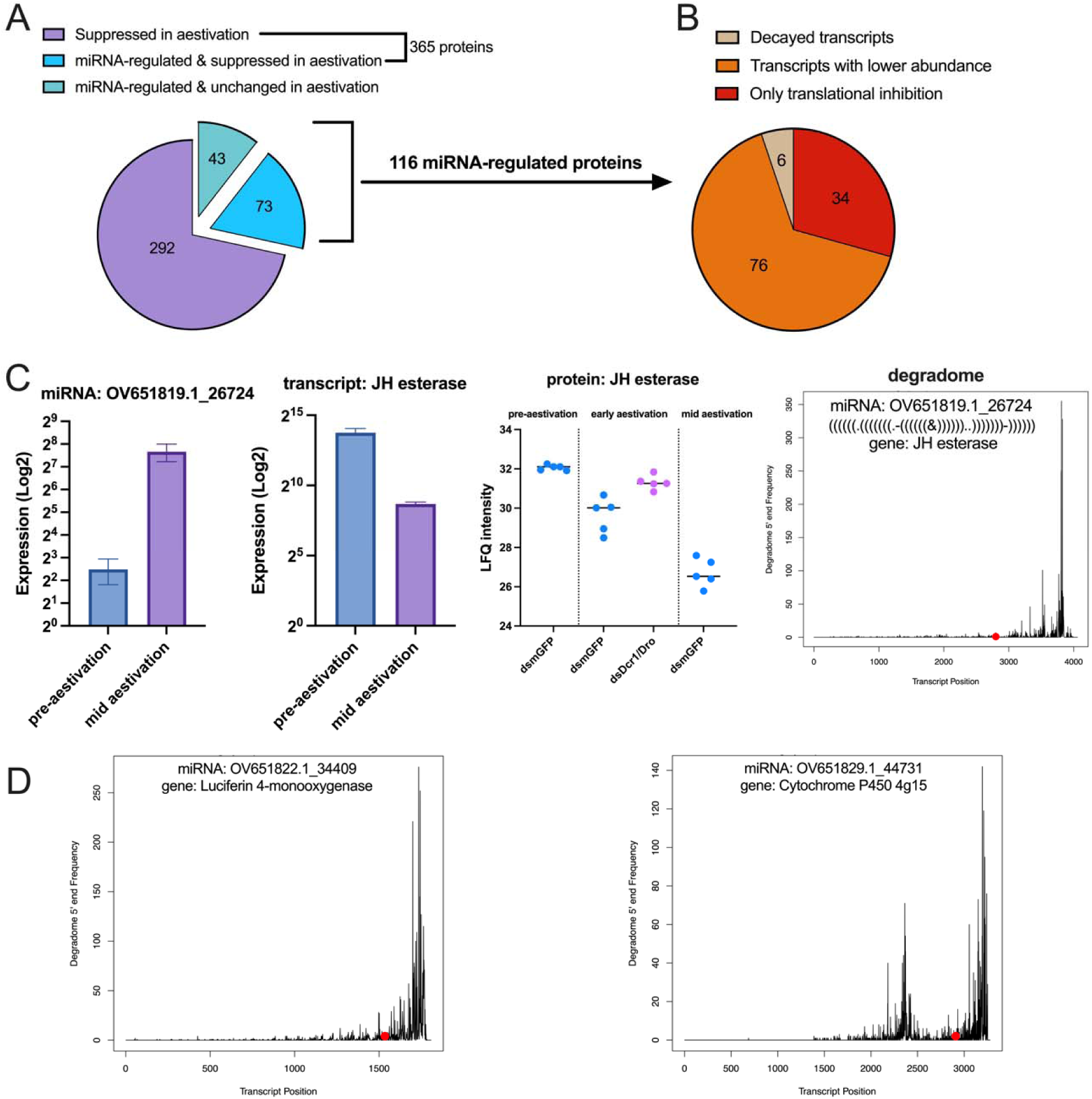
Characterizing the mode of regulation by miRNA pathway in aestivation. A) Proteomics analysis comparing aestivation and pre-aestivation in control CSFB adults identified 365 proteins with lower abundance in aestivation (Fig. S1). Out of these 365 proteins, 73 proteins were found to be deregulated upon the RNAi of miRNA pathway in aestivation (Fig. 2), while 43 proteins were also deregulated upon the inhibition of miRNA pathway in aestivation, but they did not have lower abundance in aestivation in comparison to the pre-aestivation in control CSFB. B) Among the 116 proteins that were deregulated upon the inhibition of miRNA pathway, 82 of them were found to have a lower abundance in aestivation compared to the pre-aestivation in our previous RNA-seq data (25). We only found evidence for transcript decay for 9 out of 82 of these genes. The remaining 34 proteins only had evidence for being deregulated upon the inhibition of miRNA pathway and were categorized into only translational inhibition. C) Example of a miRNA regulated gene annotated as juvenile hormone (JH) esterase. miRNA OV651819.1_26724 is overexpressed during mid aestivation (30-day-old) in intact beetles (adjusted P < 0.05, n = 3, see Fig. 1A). The transcript level of JH esterase is downregulated in mid aestivation in intact beetles (adjusted P < 0.05, n = 3, data from (25)). The protein level of JH esterase is downregulated in early (15-day-old) and mid aestivation in control beetles, while it is upregulated upon the RNAi of *dcr-1* and *dro* (dsDcr1/Dro) in early aestivation (adjusted P < 0.05, n = 5, see Fig. 2C and Fig. S1). RNA degradome suggests the deadenylation and exonucleolytic of JH esterase transcript (20 late pre-aestivation beetles were pooled into n = 1). The binding site (position 2800 nt on transcript) of complementary miRNA OV651819.1_26724 is indicated with a red dot and the complementarity between miRNA and target site is shown with dot-bracket notation. The plot was generated using CleaveLand4 and show the degradome 5’ end frequency identified on the mRNA sequence. D) Two other examples of transcripts decreasing abundance due to transcript decay mediated by miRNA increasing in abundance in mid aestivation compared to pre-aestivation.

miRNA target prediction using miRanda and 3’ UTR regions of transcripts suggested that 693 out of 3,223 transcripts with lower abundance during aestivation were targeted by at least one miRNA with higher abundance in aestivation (Fig. S2A, Dataset S3). On average, each miRNA targeted 92.93 transcripts, while each transcript was targeted by an average of 2.01 miRNAs among these predicted miRNA-transcript interactions (Fig. S2B). Focusing on transcripts encoding proteins with higher abundance upon miRNA inhibition, we found that 49 transcripts (42% of the 116 proteins) could be linked to at least one miRNA with higher abundance in aestivation through target prediction. In these cases, the average number of target transcripts per miRNA was 7.9, and each transcript was targeted by an average of 2.43 miRNAs (Fig. S2C). A similar analysis was conducted for the opposite relationship—miRNAs with lower abundance in aestivation and their predicted target transcripts with higher abundance in aestivation. This analysis identified 1,171 out of 3,013 transcripts with higher abundance during aestivation as potential miRNA targets (Fig. S2A). On average, each miRNA targeted 176.8 transcripts, and each transcript was targeted by an average of 3.3 miRNAs within these interaction (Fig. S2B).

Next, we performed transcriptome-wide RNA degradome sequencing in 10- and 15-day-old CSFB, representing late pre-aestivation and early aestivation stages, to investigate whether miRNAs with higher abundance in aestivation (Fig. 1B-C, see Dataset S1) contribute to the degradation of transcripts encoding miRNA-suppressed proteins (Fig. 2C, see Dataset S2). Analysis of degradome plots provided evidence for miRNA-mediated decay of only six transcripts (Fig. 3C-D and Fig. S3, see Dataset S4 for details). In these transcripts, we observed a high frequency of degradation at the 3’ ends, suggesting a regulatory mechanism involving deadenylation followed by exonucleolytic decay—commonly observed in animals—rather than site-specific cleavages, which is typical in plants (26) (Fig. 3C-D). Furthermore, most evidence of transcript decay was observed in the late pre-aestivation stage (10-day-old CSFB; six transcripts, Fig. S3), compared to early aestivation (15-day-old adults; two transcripts, Fig. S4), suggesting that transcript degradation predominantly occurs before aestivation. Notably, when analyzing the entire RNA degradome dataset—beyond the scope of aestivation-related genes and miRNAs—we identified four potential instances of transcript target site cleavages mediated by complementary miRNAs (categorized as "0" by CleaveLand4, Fig. S5). This finding suggests that miRNA-mediated target site cleavage may still occur in CSFB, although no such evidence was found when focusing specifically on aestivation-related regulation.

The transcripts with evidence for miRNA-mediated decay included genes that are likely important for aestivation, such as juvenile hormone (JH) esterase and cytochrome P450 (4g15)—both of which have been previously implicated in aestivation (Fig. 3C-D) (25). Together with our degradome sequencing data, we provide comprehensive evidence supporting the miRNA-mediated regulation of aestivation-related genes. For example, the miRNA OV651819.1_26724 is significantly overexpressed during mid-aestivation (30-day-old) in intact CSFB (Fig. 3C and Fig. 1A), while its complementary JH esterase transcript is downregulated during mid-aestivation in intact CSFB (adjusted P < 0.05, n = 3; data from (25)). Furthermore, at the protein level, JH esterase is downregulated during early (15-day-old) and mid-aestivation in dsmGFP-fed control CSFB, whereas it is upregulated upon dsDcr1/Dro treatment in early aestivation (adjusted P < 0.05, n = 5; Fig. 2C and Fig. S1). Additionally, RNA degradome analysis indicates decay of the JH esterase transcript. These findings collectively suggest that miRNAs play a crucial role in regulating a subset of genes involved in aestivation.

### Regulation by miRNA pathway is important for the aestivation phenotype

The finding that the inhibition of the miRNA pathway disrupts the regulation of proteins in aestivation prompted us to investigate whether the aestivation is weakened at the phenotypic level upon the inhibition of the miRNA pathway. First, we performed body composition analysis on dsDcr1/Dro-fed and control CSFB at 15 days post-emergence and 30 days post-emergence, representing early aestivation and mid-aestivation, respectively. We found that overall protein content was higher in dsDcr1/Dro-fed CSFB compared to the control group on both 15- and 30-days post-emergence (P = 0.15 and < 0.01, t = 2.88 and 7.07 by Šídák’s test, df = 28, Fig. 4A). Similarly, glucose content was higher in the dsDcr1/Dro fed CSFB, albeit only on 30 days post-emergence (P < 0.01, t = 3.9, by Šídák’s test, df = 28 Fig. 4B). In contrast, TAG levels showed an opposite pattern where dsDcr1/Dro-fed CSFB had less TAG compared to the control CSFB on both 15- and 30-days post-emergence (P < 0.01 and < 0.01, t = 6.4 and 9.5 by Šídák’s test, df = 28, Fig. 4C). We did not observe any significant changes in the trehalose and glycogen levels (P > 0.05, t < 2 by Šídák’s test, df = 28, Fig. 4D and 4E), indicating that inhibition of miRNA results in specific changes in body composition, rather than complete alteration. These results, along with the enrichment of metabolism-related proteins following miRNA pathway inhibition, suggest that miRNA is involved in maintaining a body composition appropriate for aestivation.

**Figure 4.**
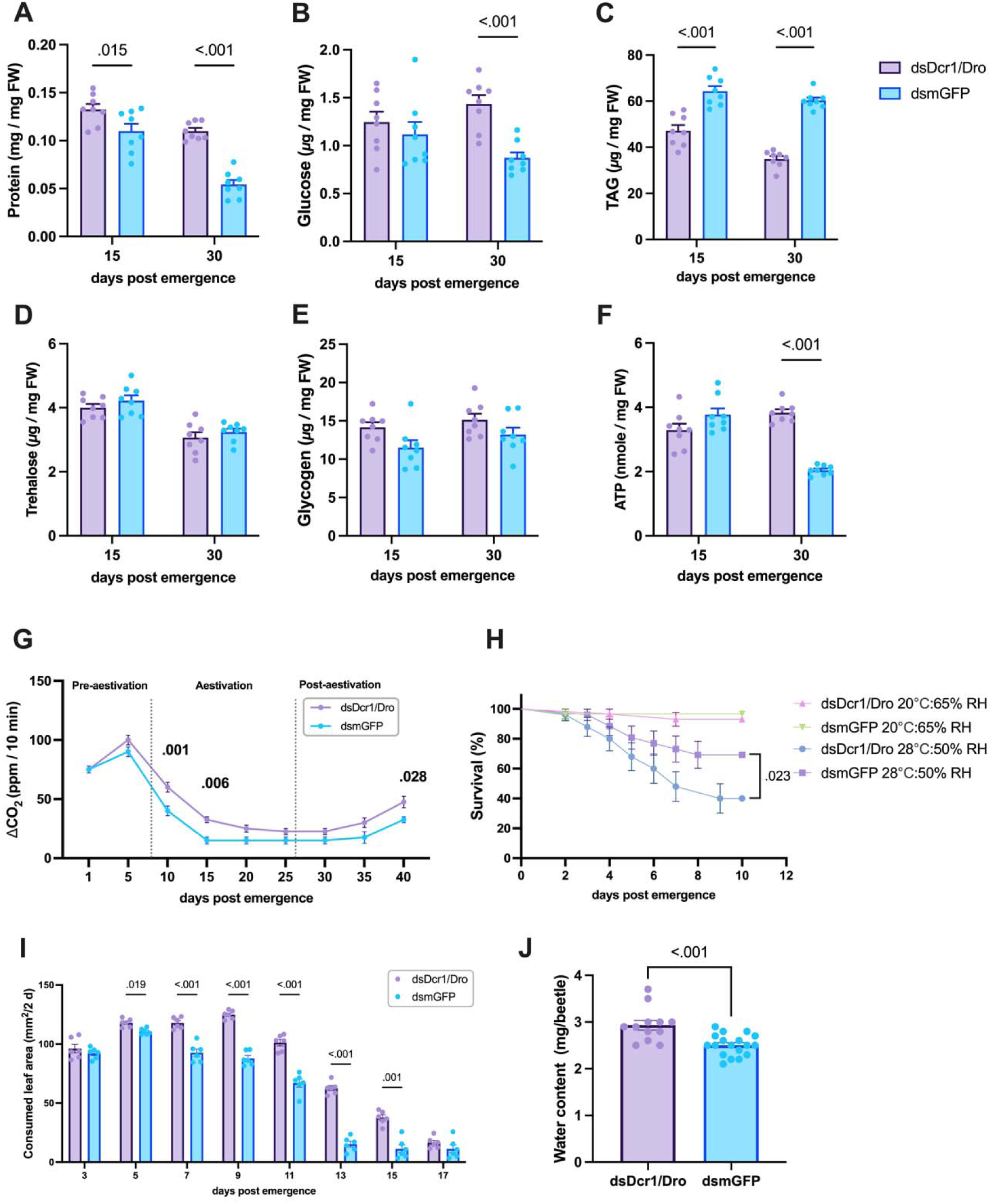
Inhibition of miRNA pathway weakens the aestivation phenotype. The miRNA pathway of CSFB adults was inhibited through feeding of chimeric dsDcr1/Dro and compared with dsmGFP fed control CSFB. Protein (A), glucose (B), TAG (C), trehalose (D), glycogen (E) and ATP (F) levels in whole bodies of 15 (early aestivation) and 30 (mid aestivation) days-old adults were measured. (G) The CO2 production was measured in CSFB post adult emergence with 5 day intervals. (H) Survival analysis with aestivating CSFB (n = 30 mixed-sex adults) under regular rearing conditions (20 °C and 65% relative humidity) or summer conditions (28 °C and 50% relative humidity). The survival of two dsRNA treatment groups receiving the same rearing condition treatment was compared using log-rank test. (I) Consumed leaf area was measured at 2-day intervals in two dsRNA treatment groups. (J) Water content measured in aestivating CSFB whole bodies and two-tailed unpaired t-test was conducted. Two-way ANOVA (factors: day and RNAi treatment) followed by Šídák’s multiple comparisons test was performed for Panels A-G and Panel I. Mean ± SEM values are provided in each panel and P < 0.05 are indicated.

It is well described that diapausing animals notably reduce their metabolic rate to maintain energy homeostasis (27). The ATP measurements showed that dsDcr1/Dro fed CSFB had significantly higher ATP content than the control group on 30-days post-emergence (P < 0.01, t = 8.3 by Šídák’s test, df = 28, Fig. 4F), but not on 15-days post-emergence (P < 0.067, t = 2.2 by Šídák’s test, df = 28). In addition, we measured CO_2_ production (VCO_2_) as a proxy for the temporal changes in metabolic rate. Notably, on both 10- and 15-days post-emergence, the dsDcr1/Dro fed CSFB had significantly higher VCO_2_ than the control CSFB (P < 0.01 and < 0.01, t = 4.1 and 3.6 by Šídák’s test, df = 54, Fig. 4G). These two time points correspond to the strong decrease in the metabolism while CSFB is entering aestivation. Although both measurements supported the idea that regulation by miRNA is important for suppressing metabolism during aestivation, the effect of miRNA inhibition on ATP seems to be delayed compared to its effect on VCO_2_.

Survival experiments were conducted under two conditions, namely 28°C temperature and 50% relative humidity (RH) and 20°C and 65% RH (laboratory rearing conditions), and initiated with 20-day-old CSFB to investigate the previously described higher resilience of aestivating beetles to hot and dry conditions (25). The dsDcr1/Dro fed CSFB had the same survival rate as the control group under laboratory rearing conditions (P = 0.55, χ2 = 0.35 by logrank test, n = 30, Fig. 4H), meaning that inhibition of miRNA does not have a detrimental effect on the survival of CSFB adults under normal circumstances. Interestingly, however, dsDcr1/Dro fed aestivating CSFB had significantly less survival than the control aestivating CSFB under hot and dry conditions (P = 0.023, χ2 = 4.03 by logrank test, n = 30, Fig. 4H). These results suggest that the regulation by the miRNA pathway is important for the resilience of aestivating CSFB to hot and dry conditions.

A defining characteristic of aestivation in CSFB, like in other insects, is the suppression of feeding activity and other activities in general. Here, we assessed whether aestivating CSFB diverged from control group in terms of feeding and activity patterns. In control CSFB, the feeding activity peaks 5 days post adult emergence and gradually declines until complete cessation of feeding activity by 15 days post-emergence (Fig. 4I). The dsDcr1/Dro fed CSFB showed higher feeding activity compared to the control CSFB following 3-days post adult emergence until 17-days post adult emergence (P < 0.05, t > 4 by Šídák’s test, df = 8). Also, the movement activity measurements showed that dsDcr1/Dro fed CSFB resumed movement activity prematurely (Fig. S6). Specifically, the miRNA-compromised CSFB moved significantly more on days 25 and 35, representing the middle and last days of the aestivation, compared to the control CSFB (P = 0.016 and 0.043, t = 3.0 and 2.7 by Šídák’s test, df = 90, Fig. S6). These results suggest that miRNA contributes to the activity inhibition in CSFB aestivation.

Water content spontaneously decreases in aestivation under normal rearing conditions in CSFB and some other insects, likely due to reduced water intake through feeding and increased dry body mass due to accumulated energy stores. Hence, we previously described the decrease in water content as an aestivation phenotype in CSFB (25). Here, water content significantly increased in dsDcr1/Dro-fed CSFB during aestivation (P < 0.01, t = 3.99, by unpaired t-test, df = 28, Fig. 4J). Hence, the increase in water content also represents a weakened aspect of the aestivation phenotype due to the inhibition of the miRNA pathway.

Overall, the inhibition of the miRNA pathway weakened the aestivation phenotype at physiological and behavioral levels, supporting the hypothesis that miRNA functionally regulates aestivation in CSFB.

## Discussion

Insect diapause involves major changes at the gene expression level, requiring dedicated gene regulatory mechanisms (28–30). Despite the discovery of key transcriptional factors required for diapause, such as FoxO (30, 31), a comprehensive picture of gene regulatory mechanisms involved in diapause is lacking. Here, we provide multifaceted evidence for the miRNA pathway being a key regulator of gene expression required for a successful aestivation (i.e., summer diapause) in the CSFB by combining small RNA-seq, mRNA-seq, proteomics, RNA degradome-seq, and functional studies through RNAi. Key factors that enabled our approach were the robust systemic RNAi response and the obligatory nature of aestivation in CSFB. Obligatory aestivation does not require manipulation of the rearing conditions to force insects into the aestivation program, meaning that the molecular readouts can be readily attributable to the aestivation program itself rather than the confounding effects of changing rearing conditions. This is important for identifying the subtle effects of miRNA on the aestivation phenotype upon the experimental inhibition of the miRNA pathway.

miRNA can regulate the abundance of a protein by causing mRNA decay and/or translational repression in animals (4, 32). Here, we found that the inhibition of the miRNA pathway mostly resulted in deregulation (increased abundance) of proteins with reduced transcript abundance in aestivation. The most parsimonious explanation is that the abundance of these proteins is regulated via mRNA decay. However, it cannot be excluded that the regulation of these proteins involved both mRNA decay and translational repression mechanisms by miRNA. Our RNA degradome analysis provided evidence for miRNA-associated decay of 6 transcripts, which was a very small proportion of all the miRNA-regulated proteins. RNA degradome analysis in animals is not as well established as in plants (33, 34). To our knowledge, comprehensive degradome data from invertebrates are available only for *Caenorhabditis elegans* (35), and we lack comparable data to define how mRNA decay occurs in beetles. Hence, further advances in RNA degradome sequencing, including both experimental and bioinformatic improvements, in non-model animals may enable further mechanistic insights and confirmation of more mRNA decay events.

In addition to demonstrating that numerous proteins are suppressed via the miRNA pathway, our RNAi experiments confirmed that miRNA-mediated regulation is essential for the majority of aestivation- associated phenotypes in CSFB. For instance, metabolism was identified as a main target of miRNA in this study, and the suppression of metabolism was compromised upon the RNAi of the miRNA pathway. Lower metabolic rates, in turn, are known to increase resilience to desiccation by allowing a reduction in spiracle activity (36, 37), explaining why miRNA-inhibited CSFB was less resilient to hot and dry rearing conditions. Likewise, another target of miRNA was digestion, and the silencing of the miRNA pathway compromised the CSFB’s ability to cease feeding activity, a major feature of aestivation and diapause in general (38–40).

Our study raises an interesting evolutionary question: Why has the miRNA gained this important function in the regulation of diapause? We argue that miRNA is an excellent fit for mediating a significant portion of the regulation required during diapause. First, usually, more genes have to be repressed rather than upregulated during diapause (25, 29, 41, 42), and repression is the dominant mode of regulation mediated by miRNA (43, 44). Second, diapausing insects often need to temporally fine-tune the expression of target genes (29, 45), which is a feature of miRNA-mediated regulation, rather than completely inhibiting their expression, which is often required for cellular differentiation and involves more robust epigenetic changes such as histone modifications (46). It should be noted that miRNA are not the only important regulators of aestivation, as only around 25% of aestivation-regulated proteins were deregulated upon the inhibition of the miRNA pathway. Rather, miRNA likely work in tandem with diapause-related transcription factors such as FoxO, which increase the transcription rate of genes involved in fat hypertrophy and survival (24). Recent studies in other insects also suggested that miRNA could be important for diapause regulation by showing differential expression of miRNA in diapausing (14, 15). However, more comprehensive studies are required in other insect species to gain evolutionary insights into the regulation of diapause through miRNA.

Climate models predict an ever-warming globe (47), and consequently increases in pest pressure on crops due to shortened generation time have already been reported (48–50). It is reasonable to argue that insect pests with pre-adaptations such as the aestivation observed in the CSFB might thrive under rising temperatures due to the existence of a genetic tool kit for becoming resilient to high temperatures (51, 52). This study showed that miRNA is a key pathway involved in regulating aestivation. Hence, studying potential changes in the miRNA pathway as temperatures rise may provide insights into whether and how these species adapt to high temperatures. This, in turn, could improve the accuracy of predicting pest species that will be more important in the future climate.

The ban on neonicotinoids and the emergence of pyrethroid-resistant populations have made the management of CSFB highly challenging, which necessitates the development of next-generation pest management strategies, such as RNA interference (53–55). The development of such strategies, in turn, requires a deep understanding of the target pest biology at the molecular level. Insights into the miRNA-mediated regulation in CSFB may support the development of innovative pest control strategies, such as those based on miRNA antagonists, to disrupt the aestivation stage of CSFB and break the pest cycle.

## Materials and Methods

### Insects

The cabbage stem flea beetles (CSFB, *Psylliodes chrysocephala*) from our laboratory colony were maintained at 20°C and 65% relative humidity under a 16:8 light/dark regime on fresh oilseed rape plants. Newly emerged adults were collected daily from pupation boxes.

### Small RNA sequencing

Total RNA was isolated from pre-aestivation (5-day old), aestivation (30-day old), and post-aestivation (55-day old) females using the Quick-RNA Tissue Kit (Zymo Research), and stored at –80°C. The same RNA samples were previously used in our mRNA-seq study (25) to ensure concordant data. Thawed RNA samples were analyzed through fragment analysis using Agilent 2100 Bioanalyzer (Agilent Technologies) to confirm integrity before library preparation. Libraries were prepared from 6 µL input RNA using the NEBNext® Multiplex Small RNA Library Prep Kit for Illumina® (NEB, catalog no: E7560S). Libraries were indexed through 13 PCR cycles, pooled by concentration, and size-selected (135–150 bp) on a 6% polyacrylamide gel. The extracted libraries were re-evaluated through fragment analysis and sequenced by BGI Tech Solutions Co. Ltd (Hong Kong) on a DNBSEQ-G400 platform (PE50+5+10 mode). Small RNA sequencing experiments were repeated four times.

### De novo miRNA prediction

The raw sRNA-seq data underwent quality analysis using FastQC (https://www.bioinformatics.babraham.ac.uk/projects/fastqc), and low-quality (q < 30) reads and adaptor sequences were removed using TrimGalore! (v0.6, https://github.com/FelixKrueger/TrimGalore). All cleaned reads were combined for de novo miRNA prediction using miRDeep2 in default mode (56). The CSFB reference genome (GenBank: GCA_927349885.1) was retrieved from NCBI for the mapper function, and mature miRNA sequences from *Tribolium castaneum* obtained from miRbase (57) acted as homologous miRNA.

### RNAi

The chimeric dsDcr1/Dro was designed to target both Pc-*dicer-1* and Pc-*drosha* using https://dsrip.uni-goettingen.de. The first 210 bp was complementary to Pc-dicer-1, while the subsequent 208 bp was complementary to Pc-*drosha* (Tab. S1). The dsDNA template for the in vitro transcription of dsDcr1/Dro and dsmGFP (control treatment (55)) were ordered as gBlocks (IDT, Germany). We used two pairs of primers, each adding T7 promoter to either end of the dsDNA template through two separate PCR reactions using Q5 high fidelity polymerase. The two strands of the dsRNA were synthesized separately using MEGAscript™ T7 Transcription Kit (Thermo Fisher Scientific, Germany) according to the manufacturer’s instructions and combined in equimolar concentration. The RNA strands were first denatured at 94 °C for 5 min and subsequently annealed at room temperature for 40 min. The length of dsRNA was confirmed through 1% agarose gel electrophoresis. dsRNA was diluted to 1 µg/µL in nuclease-free water and combined with Triton-X (200 ppm final concentration, Sigma-Aldrich). One µL of dsRNA solution was homogenously spread on 100 mm^2^ disks punched out from young oilseed rape leaves. Freshly treated leaf disks were provided to newly emerged CSFB adults every 2 days until 17 days post-adult emergence.

### Proteomics

Whole bodies of 4 female CSFB treated with either dsDcr1/Dro or dsmGFP were collected at 5, 15, or 30 days post-adult emergence, flash-frozen in liquid nitrogen and homogenized by grinding. Proteins from the samples were extracted using urea denaturation buffer, which contained 6 M urea, 2 M thiourea and 10 mM HEPES (pH 8.0) at a proportion of 1 ml per 100 mg homogenized sample. The supernatant containing protein was recovered through centrifugation at 12,000 g for 5 min. Bicinchoninic acid assay (BCA) was used to determine protein concentration. Next, proteins were precipitated using chloroform/methanol extraction (58). The protein pellet was suspended in Rapigest SF solution (0.1%, Waters) and digested using trypsin (1:20, Serva, Germany). The digested protein was prepared for LC/MS analysis by adding TFA at a 1:10 ratio, incubating the mixture at 37 °C for 45 min, and subsequently drying it using a SpeedVac. A standard C18 stage tipping protocol was performed to purify the peptides (59, 60). The proteomics experiments were repeated for a total of 5 biological replicates.

LC/MS analysis of the peptide samples was performed with an Ultimate 3000 system (Thermo Fisher Scientific) coupled to a Q Exactive HF mass spectrometer (Thermo Fisher Scientific). Peptides were separated by reversed phase liquid chromatography and on-line ionized by nano-electrospray (nESI) using the Nanospray Flex Ion Source (Thermo Fisher Scientific). Full scans were recorded in a mass range of 300 to 1650 m/z at a resolution of 30,000. Data-dependent top 10 HCD fragmentation was performed at a resolution of 15,000 with dynamic exclusion enabled. The XCalibur 4.0 software (Thermo Fisher Scientific) was used for LC-MS method programming and data acquisition. MaxQuant (v2.6) and Perseus (v1.6) proteomics software packages were used for data analysis. Calculation of label-free quantification values (LFQ) was performed, and a CSFB-adult-specific protein database was used that was generated from our transcriptome data (25). Proteins with fewer than three peptide counts were discarded. Proteins with an adjusted P < 0.05 and either log_2_ fold > 1 or log_2_ fold < −1 were considered significantly increased or decreased in abundance, respectively.

### miRNA target prediction

The target transcripts of miRNA showing differential abundance in aestivation were predicted using miRanda using the "strict" mode and 140 as miRanda score treshold. As the target input, the 3’ UTR regions from the direction corrected CSFB transcriptome was extracted using TransDecoder (https://github.com/TransDecoder/TransDecoder/wiki). Next, the transcripts showing differential abundance in aestivation (25) were identified.

### RNA degradome sequencing

We isolated total RNA from 20 intact 10- or 15-day-old CSFB female whole bodies, using the Quick-RNA Tissue kit (Zymo Research). RNA quality was assessed through fragment analysis using Agilent 2100 Bioanalyzer (Agilent Technologies), and 100 µg of RNA was used as input for degradome library preparation based on a published method (61). Polyadenylated RNA was purified using Dynabeads mRNA Purification kit (ThermoFisher) and Oligo d(T) 25 Magnetic Beads (NEB). The 5’ RNA adaptor (100 µM, Tab. S2) was ligated to RNA using T4 RNA ligase buffer (NEB) and 5’ RNA adaptor ligated polyadenylated RNA was purified using Dynabeads mRNA purification kit (ThermoFisher). First, the cDNA strand was synthesized using ProtoScript TM II reverse transcriptase (NEB). The reaction for the second strand cDNA synthesis included first cDNA strand, MyFi Mix (BioCat), and 5’ and 3’ adaptor primers (Tab. S2), and 15 PCR cycles were performed. The PCR product was purified using Monarch PCR & DNA cleanup kit and digested with EcoP151 (NEB). dsDNA adaptors (Tab. S2) were ligated to the digested DNA using T4 DNA ligase (NEB). The dsDNA adaptor-ligated DNA (79 bp) was purified through PAGE and concentrated through ethanol precipitation. Thirteen PCR cycles were performed using MyFi Mix (BioCat) to add the SR1 indexes (Tab. S2) to the libraries. The libraries were pooled and purified through PAGE. The sequencing service was provided by BGI Tech Solutions Co. Ltd (Hong Kong) on a DNBSEQ-G400 platform (PE50+5+10 mode).

### RT-qPCR and sRNA composition

Four CSFB females receiving dsDcr1/Dro or dsmGFP treatment were flash-frozen in liquid nitrogen after wing removal. Total RNA from CSFB samples was extracted using the Quick-RNA Tissue & Insect MicroPrep™ Kit (Zymo Research, Germany). RT-qPCR was conducted with the Luna One-step RT-qPCR kit (New England Biolabs, Germany) on a 384-well plate, with three technical replicates per biological replicate, using the CFX384 Touch Real-Time PCR system (Bio-Rad). Each well contained 0.8 μL of total RNA (500 ng/μL) and 0.8 μL of a 10 mM forward and reverse primer mix (IDT) to measure the expression of *Pc*-*dicer-1* and *Pc*-*drosha* and a reference gene, *Pc*-*rps4e*, optimized for RNAi experiments by Cedden et al. (2024) (primers in Tab. S2). The normalized expression values were obtained using CFX Maestro Software (Bio-Rad).

The total RNA extracted from 15-day-old CSFB adults were subjected to fragment analysis using Agilent 2100 Bioanalyzer (Agilent Technologies), and the miRNA percentage was calculated using the ProSize data analysis software. These experiments were repeated three times.

### Body composition

The fresh weights of individual CSFB were measured to normalize the following body content measurements. The whole bodies were frozen using liquid nitrogen and homogenized by grinding. Trehalose from the samples was measured using the Trehalose Assay Kit (Megazyme). Glucose was measured using the D-Glucose HK Assay kit (Megazyme) and separated homogenized samples were first incubated with 0.5 units of amyloglycosidase for 4 hours at 37°C to obtain glycogen content. Triglyceride content was obtained by calculating the difference between total glycerol and free glycerol for each sample using Triglyceride Colorimetric Assay Kit (Cayman) and Free Glycerol Reagent Kit (Sigma), respectively. The ATP content was measured using ATP Assay Kit (Sigma-Alrdrich). Water content was measured by calculating the difference between initial fresh weight and weight after drying at 60°C for 72 h. Body composition analyses were repeated 8 times.

### Survival analysis

CSFB adults receiving dsDcr1/Dro or dsmGFP (n = 30 mixed-sex adults that were 20-day-old at the start of the experiment) were transferred into climate chambers set to 28 °C and 50% relative humidity or regular rearing conditions (20 °C and 65% relative humidity). The survival of individual beetles was monitored daily using a fine brush, and those showing no movement were considered dead.

### Feeding and movement activity

The leaves were flattened beside a 1 cm reference line for calibration and photographed using a 25-megapixel camera. The remaining leaf area was calculated with ImageJ software (v1.8, https://imagej.nih.gov) using the ‘Analyze Particles’ function, and this area was subtracted from the initial leaf disk area (400 mm²).

Beetles were individually placed in vented Petri dishes (100□mm × 15□mm) and tracked using a Zantiks LT unit (Zantiks Ltd.) with a 4 ×□5 layout under white light at room temperature. Movement was recorded in 5□s bins over 2□h. Total movement (mm) was summed per beetle, and non-zero bins were averaged to calculate speed (mm/s). The script for this measurement is available on FigShare: https://doi.org/10.6084/m9.figshare.24162525.v1.

### Statistical analysis

Cleaned sRNA reads were used to calculate raw count values of mature miRNA sequences per library using miRDeep2’s "quantifier" function. These raw counts served as input for differential expression analysis with DESeq2 (v1.46) (62). Aestivation vs. pre-aestivation and aestivation vs. post-aestivation comparisons were conducted, and miRNA with P < 0.05 were accepted as differentially abundant.

Proteins with fewer than three peptide counts were discarded. Proteins with an adjusted P < 0.05 and either log_2_ fold > 1 or log_2_ fold < −1 were considered significantly increased or decreased in abundance, respectively (n = 5). The proteins with significantly higher abundance were subjected to GO enrichment analysis using R package "clusterProfiler" (v3.17) and our previous functional annotation for CSFB adults generated by Trinotate (v4.0, https://github.com/Trinotate) (25). The most significantly enriched 15 GO pathways were plotted.

The normalized expression data for each target gene was obtained from RT-qPCR were analyzed using Two-way ANOVA (factors: day and treatment) followed by Šídák’s multiple comparisons test to compare the RNAi and control groups (n = 3). miRNA percentage values were analyzed through unpaired one-tailed t-test (n = 3).

The degradome frequency plots were generated by mapping degradome reads to our CSFB adult transcriptome using CleaveLand4 (v4.5, https://github.com/MikeAxtell/CleaveLand4). The degradome frequency plots were manually investigated to find evidence for decay of mRNA that were complementary to miRNA with higher abundance in aestivation.

Survival data was analyzed using logrank test to compare the RNAi and control groups reared under different conditions (n = 30 per group). Body composition (n = 8), VCO_2_, (n = 4, paired) feeding (n = 6, paired), and movement (n = 10) data were analyzed using two-way ANOVA (factors: day and treatment) followed by Šídák’s multiple comparisons test to compare the RNAi and control groups. Water content data was analyzed using two-tailed unpaired t-test.

## Acknowledgments

The authors D.C. and G.G. were funded by the Deutscher Akademischer Austauschdienst (DAAD) research grants program. The authors thank Dr. Franziska Beran and Daniel Veit for providing the modified LI-820 CO_2_ Gas Analyzer. The authors also thank Dr. Oliver Valerius, Dr. Miki Kawachi-Reuscher, and Katharina Ziese-Kubon for supporting the experiments.

## Author Contributions

G.G. and D.C. conceptualized and designed research; G.G., K.S., O.V., J.Z., U.T., M.R., S.S., and DC contributed new reagents/analytic tools; G.G. and D.C. performed research and analyzed data; D.C. supervised the research; M.R., and S.S. supported research; G.G. and D.C. wrote the manuscript. G.G., U.T., M.R., S.S., D.C. revised and edited the manuscript.

