## Supporting information for "The MicroRNA pathway regulates obligatory aestivation in a flea beetle"

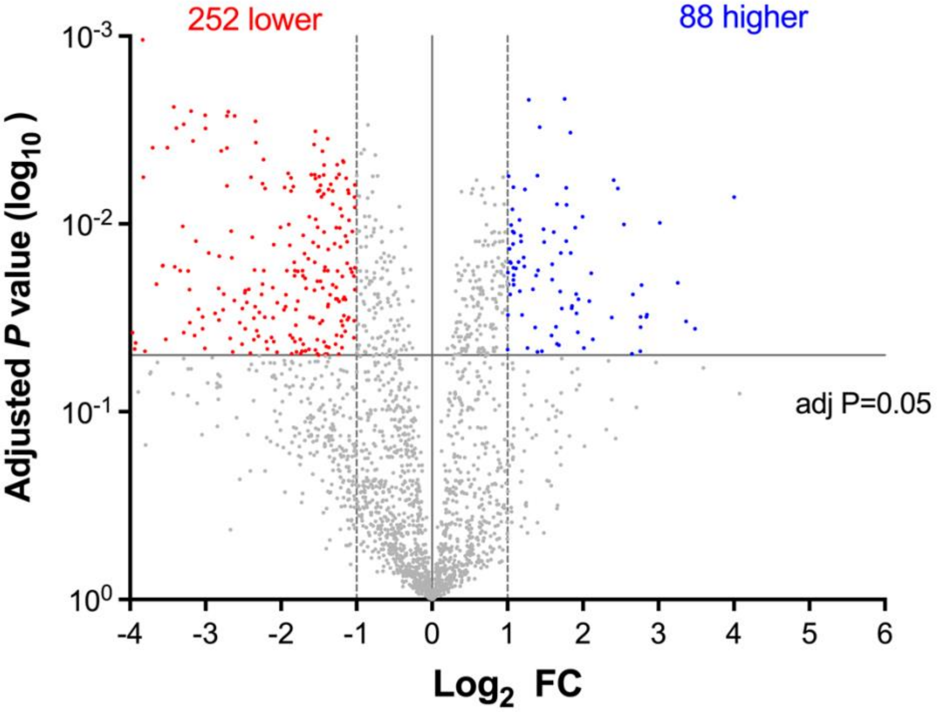


**Figure S1. Differentially abundant proteins in aestivation (15-day-old) compared to pre-aestivation (5-day-old) dsmGFP-fed (control) cabbage stem flea beetle (n = 5).**

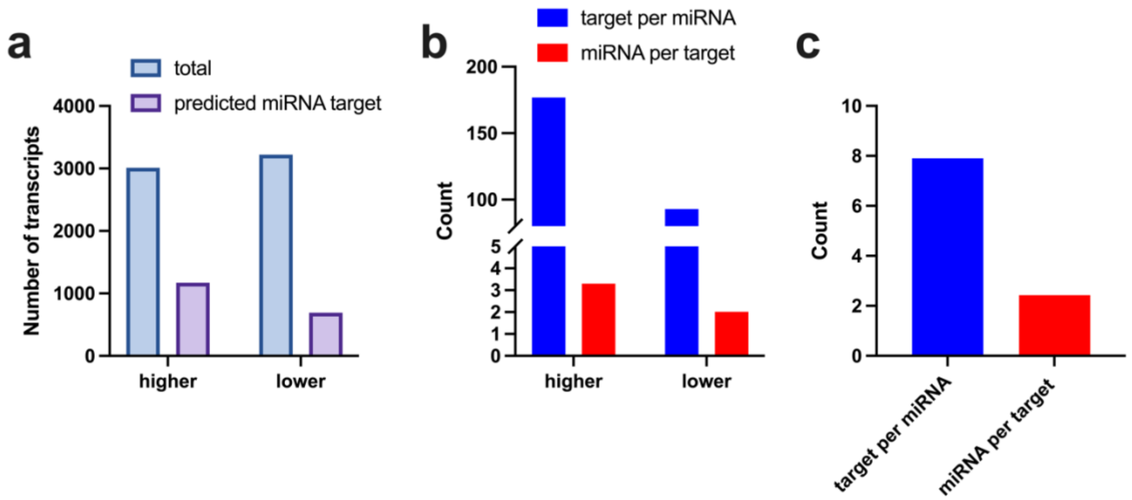


**Figure S2. Integration of miRanda predictions for miRNA differentially abundant in aestivation with transcripts differentially abundant in aestivation that showed reverse expression.** a) Number of total transcripts with higher or lower abundance in aestivation and the subset of these predicted to be targeted by at least one miRNA with reverse expression in aestivation. b) Counts of target transcripts per miRNA and miRNA per target transcripts among the  transcript with higher or lower abundance in aestivation. b) Counts of target transcripts per miRNA and miRNA per target transcripts among the transcript with  lower abundance in aestivation in addition to encoding for a protein determined to be miRNA regulated through RNAi.

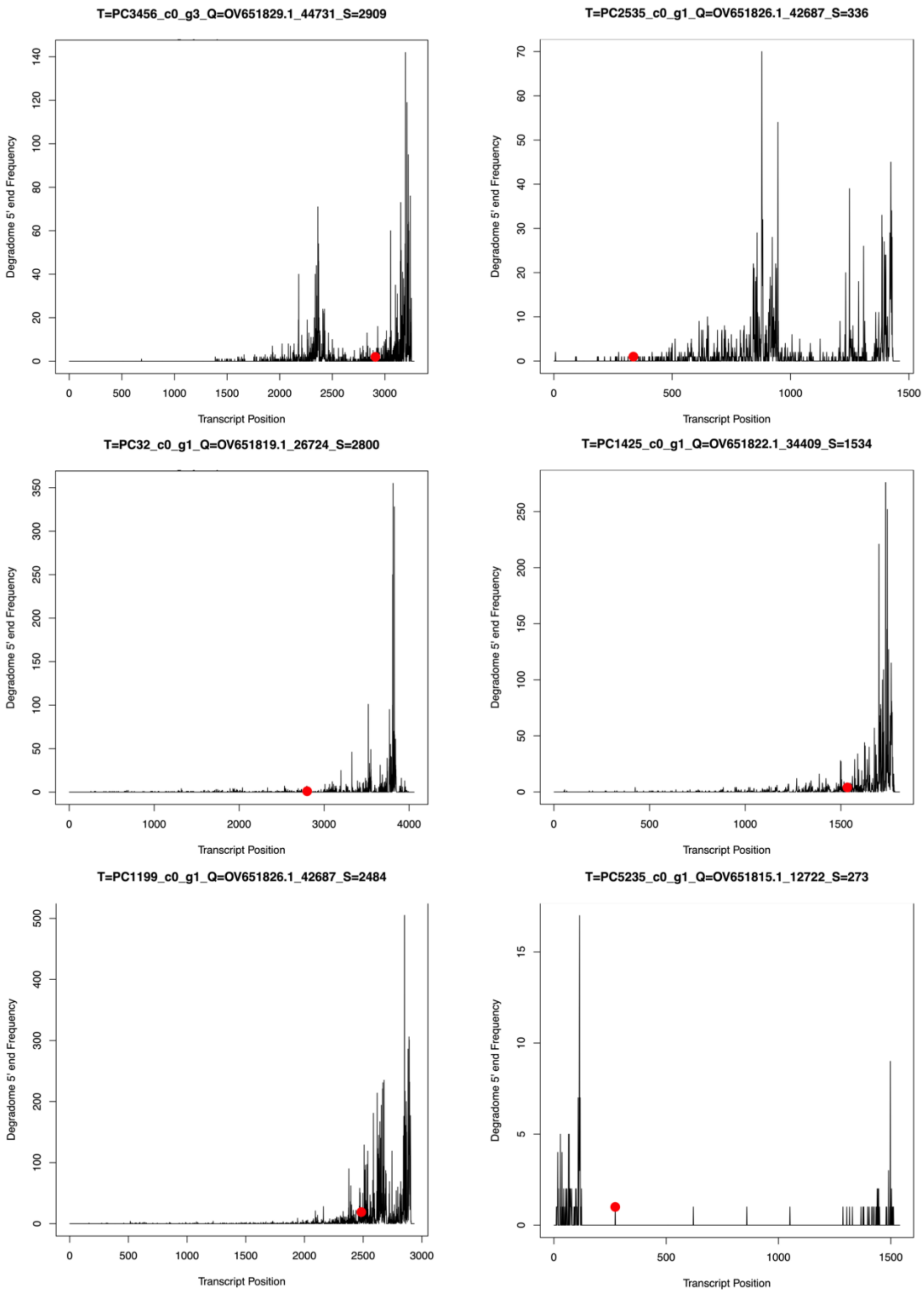


**Figure S3. Evidence for transcript decay mediated by miRNA with higher abundance in aestivation.** RNA degradomics was conducted in 10-day-old cabbage stem flea beetle adults. Twenty whole bodies were pooled into one sequencing lane. The region complementary to the miRNA is indicated with a red dot. RNA degradome suggests the exonucleolytic decay of the transcripts.

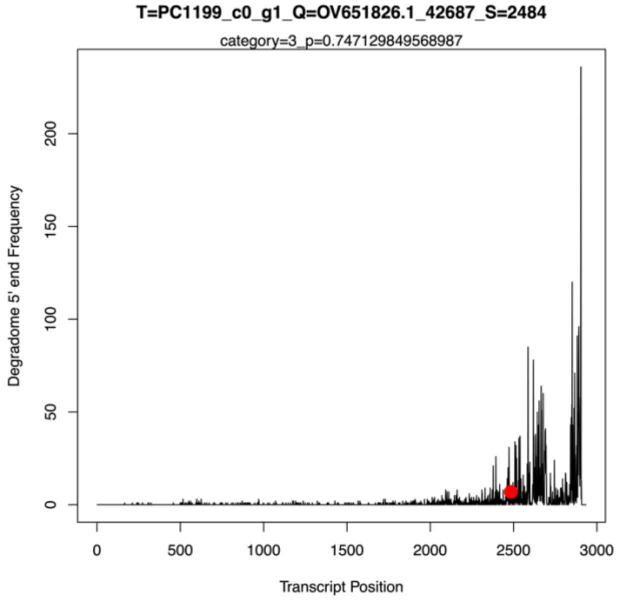


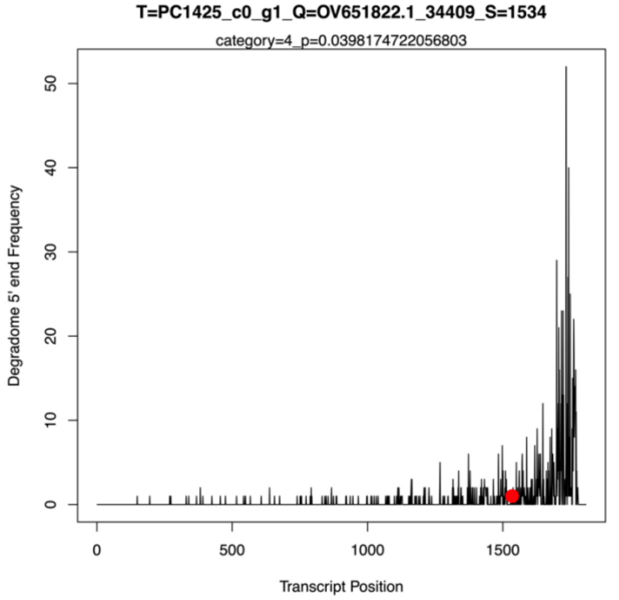


**Figure S4. Evidence for transcript decay mediated by miRNA with higher abundance in aestivation.** RNA degradomics was conducted in 15-day-old cabbage stem flea beetle adults. Twenty whole bodies were pooled into one sequencing lane. The region complementary to the miRNA is indicated with a red dot. RNA degradome suggests the exonucleolytic decay of the transcripts.

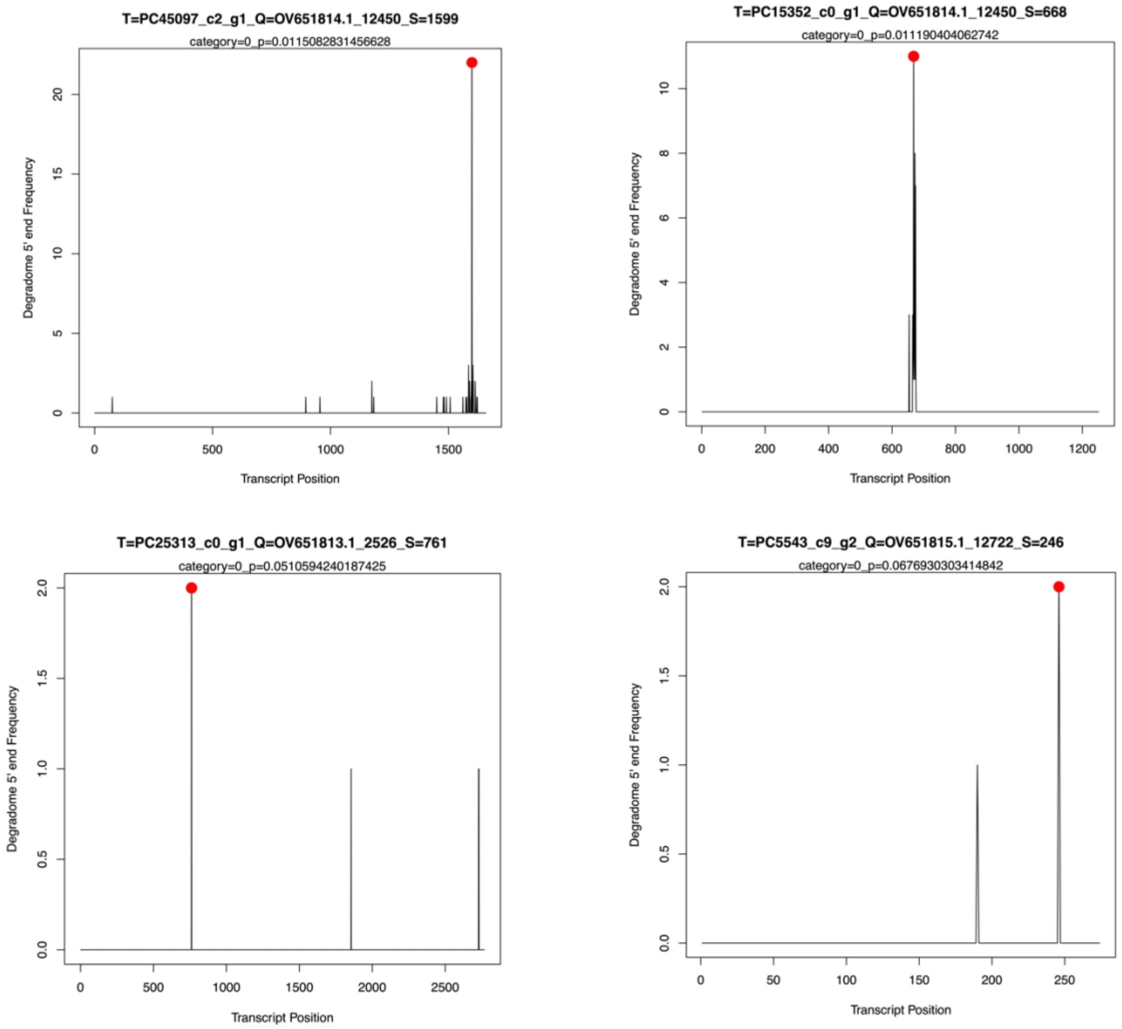


**Figure S5. Evidence for target site cleavages mediated by miRNA in the transcriptome of cabbage stem flea beetle.** Plots were generated by CleaveLand4 and they were categorized as “0” meaning highest evidence for target site cleavage. The region complementary to the miRNA is indicated with a red dot. However, these transcripts or miRNA were not found to be related to aestivation in our other datasets (see Fig. 1 and Fig. 2 in the main manuscript). These suggest that target site cleavages mediated by miRNA may rarely take place in CSFB.

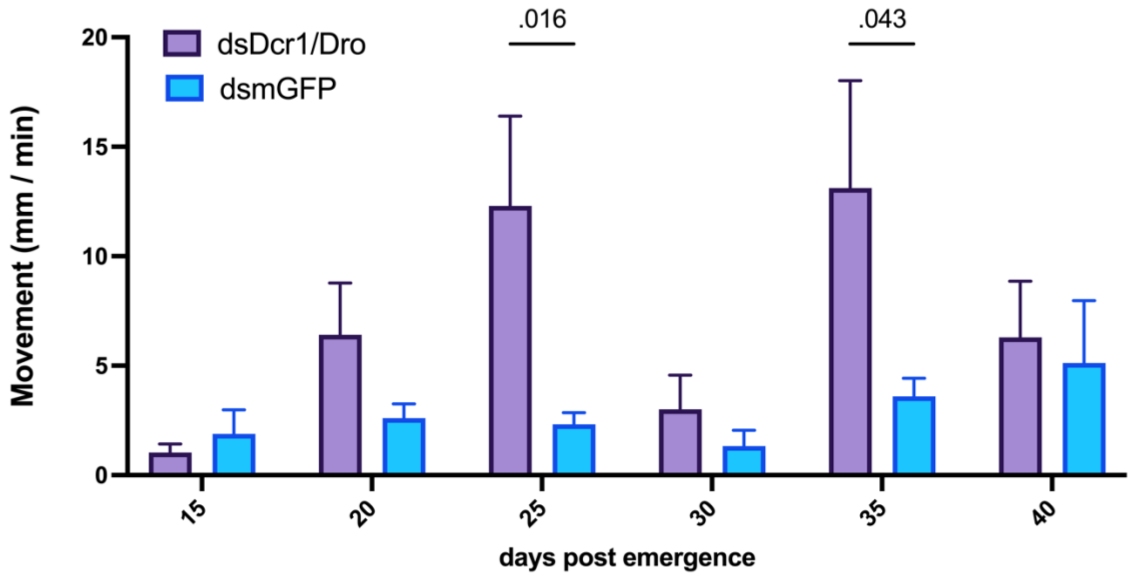


**Figure S6. Temporal movement activity measurements in CSFB adults fed with dsDcr1/Dro or dsmGFP (n = 10 per treatment).** The movement activity was measured using Zantiks LT unit for 2 h spanning the morning activity peak and statistically analyzed using two-way ANOVA followed by Šídák's multiple comparison test.

**Table S1. Sequence of chimeric dsDcr-1/Dro**

| |
|---|
| ATCCAAGCTCATGTTCCTGGTAGCAGCCATGTCAGCGACTGACCAACAATTAGCGGAAAT<br>TGAGCATCGATAAGAGCTTCCTCTAGTTGTTTGGGTACGTAAAAACATGGTGGTAACCAGT<br>TGTCGTGAGGATCAAATTTTGTAGCTATCATAAACTCGCCCAGATTTTTGCGCCTGCCCAAT<br>CGATATAAGTTCAAATTGCTGACTTGGCCTTTTTGGCGGCGTTCATTTCGGCTTGCTGGATA<br>CTGTGGCCCATTGCACTAGCCAGTCTCCTACCTCTAAAATACACAGCTACTGTATACACTCT<br>CGTATTAGTTGGTCCTTTACATTCAATGACCTTATAAACCGGAATATCGGGTTCGCCGCCGT<br>CCATAGTACGCAAAGTAAGGCAGCACTGCTGCAATTTCGATTTGGGAT |

*The first 210 base pairs of the dsRNA targets Pc-*dicer-1* (yellow) and the subsequent 208 base pairs target Pc-*drosha* (aqua). The transcriptome for CSFB adult is available on NCBI (BioProject: PRJNA930726 and TSA: GKIH00000000.1). The accessions for Pc-*dicer-1* and Pc-*drosha* are GKIH01052927.1 and GKIH01081364.1, respectively.

### Table S2. Sequences of the primers and adaptors used in the study

| Name | Purpose | Direction | Sequence |
|---|---|---|---|
| Pc-*dicer-1*[1] | RT_qPCR | Forward | TCCCGATGATCAACGTAGCG |
| Pc-*dicer-1*[1] | RT_qPCR | Reverse | TATTCGGACCCAGGGAATGC |
| Pc-*drosha*[2] | RT_qPCR | Forward | ACGCAGACGAAATCAAGGGT |
| Pc-*drosha2* | RT_qPCR | Reverse | TGCGGTGGTCTGATTCCAAA |
| dsmGFP | In vitro transcription | Forward | GGACCCTGACCTACGGCCTAT |
| dsmGFP | In vitro transcription | Reverse | GGTGCCGTCCTCGTACTAGTT |
| dsmGFP | In vitro transcription | Forward with T7 | GAATTGTAATACGACTCACTATAGGACCCTGACCTACGGCCTAT |
| dsmGFP | In vitro transcription | Reverse with T7 | GAATTGTAATACGACTCACTATAGGTGCCGTCCTCGTACTAGTT |
| dsDcr-1/Dro | In vitro transcription | Forward | ATCCAAGCTCATGTTCCTGGTAGC |
| dsDcr-1/Dro | In vitro transcription | Reverse | ATCCCAAATCGAAATTGCAGCAG |
| dsDcr-1/Dro | In vitro transcription | Forward with T7 | TAATACGACTCACTATAGGATCCAAGCTCATGTTCCTGGTAGC |
| dsDcr-1/Dro | In vitro transcription | Reverse with T7 | TAATACGACTCACTATAGGATCCCAAATCGAAATTGCAGCAG |
| *Pc-rps4e*[3] | RT_qPCR | Forward | GGGTCGTGTTGGTACGGTAA |
| Pc-*rps4e*[3] | RT_qPCR | Reverse | AGTAGCAAACACGTGGCCAT |
| 5′ RNA adaptor | RNA degradome | Forward | GUUCAGAGUUCUACAGUCCGACGAUCAGCAG |
| RT-primer | RNA degradome | Reverse | CGAGCACAGAATTAATACGACTTTTTTTTTTTTTTTTTT |
| 5′ adaptor | RNA degradome | Forward | GTTCAGAGTTCTACAGTCCGAC |
| 3′ adaptor | RNA degradome | Reverse | CGAGCACAGAATTAATACGACT |
| dsDNA top | RNA degradome | Forward | NNTGGAATTCTCGGGTGCCAAGG |
| dsDNA bottom | RNA degradome | Reverse | CCTTGGCACCCGAGAATTCCA |
| Final 5′PCR primer | RNA degradome | Forward | AATGATACGGCGACCACCGAGATCTACACGTTCAGAGTTCTACAGTCCGA |
| Final 3′PCR primer for 10d-old CSFB | RNA degradome | Reverse with index | CAAGCAGAAGACGGCATACGAGATTTAGGCGTGACTGGAGTTCAGACGTGTGCTCTTCCGATCT |
| Final 3′PCR primer for 15d-old CSFB | RNA degradome | Reverse with index | CAAGCAGAAGACGGCATACGAGATGATCAGGTGACTGGAGTTCAGACGTGTGCTCTTCCGATCT |

1Accession for Pc-dicer-1 is GKIH01052927.1 in TSA for CSFB adult: GKIH00000000.1

2Accession for Pc-drosha is GKIH01081364.1 in TSA for CSFB adult: GKIH00000000.1

3Accession for *Pc-rps4e* is GKIH01045819.1 in TSA for CSFB adult: GKIH00000000.1

**Dataset S1 (separate file).** De novo identification of and differential abundance analysis on miRNA at different life stages in the cabbage stem flea beetle

**Dataset S2 (separate file).** Differential abundance analysis on proteins following the inhibition of miRNA pathway

**Dataset S3 (separate file).** miRNA target prediction by miRanda

**Dataset S4 (separate file).** RNA degradomics analysis in 10- and 15-day-old cabbage stem flea beetle adults
